# The aging DNA methylome reveals environment-by-aging interactions in a model teleost

**DOI:** 10.1101/2021.03.01.433371

**Authors:** Emily M. Bertucci, Marilyn W. Mason, Olin E. Rhodes, Benjamin B. Parrott

**Affiliations:** Odum School of Ecology, University of Georgia, Athens, GA, 30602, USA; Savannah River Ecology Laboratory, University of Georgia, Aiken, SC, 29802, USA

## Abstract

The rate at which individuals age underlies variation in life history and attendant health and disease trajectories. Age specific patterning of the DNA methylome (“epigenetic aging”) is strongly correlated with chronological age in humans and can be modeled to produce epigenetic age predictors. However, epigenetic age estimates vary among individuals of the same age, and this mismatch is correlated to the onset of age-related disease and all-cause mortality. Yet, the origins of epigenetic-to-chronological age discordance are not resolved. In an effort to develop a tractable model in which environmental drivers of epigenetic aging can be assessed, we investigate the relationship between aging and DNA methylation in a small teleost, medaka (*Oryzias latipes*). We find that age-associated DNA methylation patterning occurs broadly across the genome, with the majority of age-related changes occurring during early life. By modeling the stereotypical nature of age-associated DNA methylation dynamics, we built an epigenetic clock, which predicts chronological age with a mean error of 29.1 days (~4% of average lifespan). Characterization of clock loci suggests that aspects of epigenetic aging are functionally similar across vertebrates. To understand how environmental factors interact with epigenetic aging, we exposed medaka to four doses of ionizing radiation for seven weeks, hypothesizing that exposure to such an environmental stressor would accelerate epigenetic aging. While the epigenetic clock was not significantly affected, radiation exposure accelerated and decelerated patterns of normal epigenetic aging, with radiation-induced epigenetic alterations enriched at loci that become hypermethylated with age. Together, our findings advance ongoing research attempting to elucidate the functional role of DNA methylation in integrating environmental factors into the rate of biological aging.

## Introduction

Selective pressures resulting from environmental conditions modify the rate of aging over time and are hypothesized to contribute to variation in lifespan across species (1). At the individual level, variation in the rate of aging reflects a decoupling of biological from chronological aging which may underlie variable timing of life history events, physiological function, and the onset of age-related disease (2,3). While the origins of biological-to-chronological age mismatch are unknown, evidence suggests that the environment is a key determinant (2,4). However, the mechanisms translating environment attributes into variation in biological aging are not fully resolved.

One potential mechanism integrating environmental factors into variable aging trajectories is age associated patterning of the epigenome. DNA methylation represents a fundamental epigenetic modification and occurs in concert with many other epigenetic processes, including histone modifications (5). Not only is variation in the DNA methylome linked to extrinsic factors such as exposure to contaminants (6), famine (7), season (8) and social environment (9), recent studies show compelling links between epigenetic changes and aging (2,10,11). Stereotypical changes occurring in the DNA methylome with age can be modeled to construct “epigenetic clocks” which predict chronological age with unprecedented accuracy in several vertebrate species (2,12). While epigenetic modifications are considered a hallmark of normal aging (13), the causal role “epigenetic aging” might play in declines in cellular and physiological function is debated. Additional research on the basic biology of epigenetic aging in a variety of species is needed to advance a broader understanding of the underlying mechanisms driving both normal and accelerated aging.

Certain aspects of epigenetic aging appear conserved across vertebrates, with epigenetic clocks developed in several mammals (10,14–17), a bird (18), and most recently fish (19,20). Comparisons of epigenetic clocks in mice, humans, and dogs reveal functional similarities including an enrichment of age-associated loci in developmental genes (21), suggesting these clocks might converge on similar biological processes across taxa. The heritability of epigenetic age acceleration has been estimated to be up to 0.4 (10), although more recent evidence suggests this is an overestimation and that shared environmental conditions exert more influence on epigenetic age acceleration when compared to genetic components (22). Thus, whereas phylogenetic relationships may play a role in determining the rate of epigenetic aging across species, environmental conditions are likely the most significant drivers of intraspecific variation in epigenetic aging.

Epigenetic age acceleration in humans is correlated with environmental factors including stress (23) and pollution (24), and in mice associates with environmental conditions known to extend lifespan, such as caloric restriction (25,26). Interestingly, epigenetic age acceleration in humans is also connected to precocious timing of key life history traits including the onset of puberty (27), menopause (28), and mortality (29). Even more, epigenetic age appears to reflect aspects of reproductive effort, such as oocyte yield (30), number of pregnancies (31), and the birth weight of offspring (32). Given that adverse environments are also thought to affect plasticity in life history traits (1), epigenetic aging presents a potential mechanism linking environmental factors and individual variation in biological aging. However, identifying causal relationships between epigenetic aging and environmental factors is fundamental to understanding the mechanistic role epigenetic processes might play in life history variation.

Alterations to the epigenome are recognized as a hallmark of aging (13,33,34), but whether age-associated epigenetic modifications are reflective of one or many different underlying processes is unknown. Some of the first studies reporting epigenetic change with age reported the random loss of epigenetic information over time, or epigenetic drift (35). In contrast, epigenetic clocks rely on non-random changes to the DNA methylome over time, suggestive of a process independent of epigenetic drift (36). Recent work demonstrates that DNA damage, namely the induction of DNA double stranded breaks (DSBs), plays an important role in the acceleration of epigenetic age (37). Induction of DSBs in mice accelerates epigenetic aging via the relocalization of chromatin modifiers (RCM) which are recruited to aid in DSB repair (37). During RCM, chromatin modifiers leave their original loci and do not always return with high fidelity, resulting in aberrant epigenetic patterning and the loss of epigenetic information over time (37). Interestingly, this proposed mechanism involves the occurrence of epigenetic drift at non-random loci (37) – producing predictable patterns from random processes. This provides a potential link between the endogenous and exogenous factors which may accelerate normal aging (38). However, additional research is warranted to understand whether epigenetic aging and epigenetic drift are separate, indistinguishable, or linked processes. Further, resolving the relationship between environmental factors (i.e., those causing DSBs) is required to understand the origins of epigenetic-to-chronological age discordance and the impact it may have on variation in life history traits.

Here, we investigate signatures of epigenetic aging by sequencing DNA methylomes of a short-lived teleost, the medaka fish (*Oryzias latipes*), and compare our findings to those observed in mammals. We then construct an epigenetic clock *de novo* based on age-associated changes to the DNA methylome and use an outdoor replicated mesocosm experiment to analyze how chronic exposure to ionizing radiation (IR), an environmental factor known to cause DSBs, affects normal epigenetic aging. While associations between epigenetic age acceleration and disease states are well established, the proximate mechanisms regarding how epigenetic aging interacts with environmental signals is not known. To address this, we investigate the physical convergence of environmentally induced changes and age-related changes across the DNA methylome. Collectively, our study advances our understanding of conserved epigenetic aging patterns in vertebrates, and highlights how environmental factors, like IR, influence the aging methylome to both disrupt and accelerate normal epigenetic aging.

## Results

### Characteristics of the age-associated methylome

Spearman correlation coefficients revealed that 1550 loci (1.95%) gain methylation with age (cor > 0.5) and 604 loci (0.76%) lose methylation with age (cor < −0.50; Figure 1A). Collectively, 2.7% of all loci covered were correlated with age as assessed by correlation coefficients > ± 0.5. After an FDR correction for multiple comparisons, 62 loci (0.08%) retained significant p-values (*p* < 0.05). Thus, age-associated loci are generally hypomethylated in young fish (2-months) relative to older fish (6- and 12-months) and gain methylation with age (Figure 1B).

**Figure 1.**
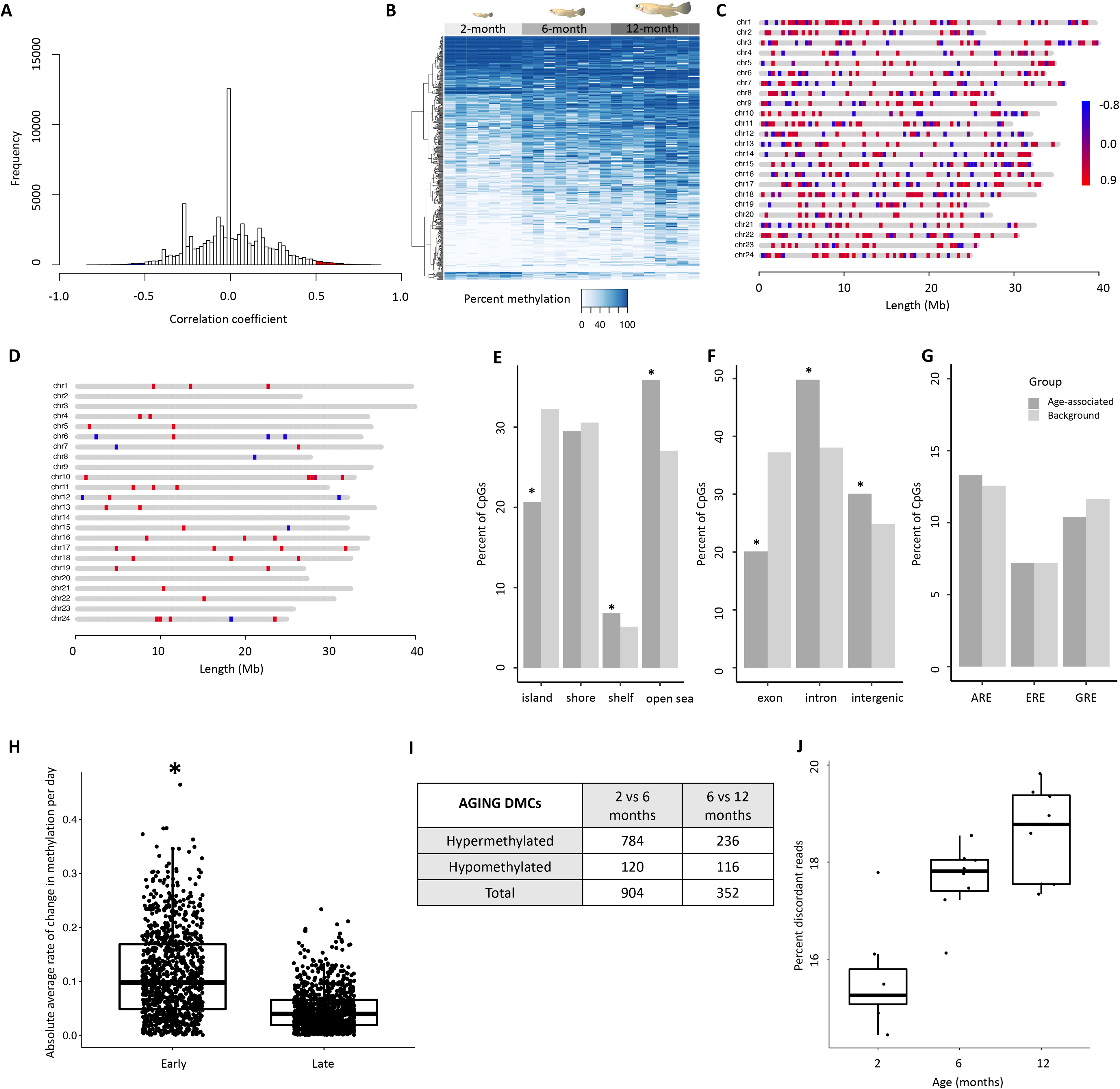
Characterization of age-associated DNA methylation patterning in medaka hepatic tissue. (A) Histogram of correlation coefficients between methylation status and age in days. Hypermethylated loci are shown in red and hypomethylated in blue. (B) Heatmap of the top 500 age-associated loci. Age is specified by color (2-month: light gray, 6-month: medium gray, 12-month: dark gray). (C) Distribution of the top 1000 age-associated loci and 62 significantly associated loci (D) across the medaka genome. Loci that become hypermethylated with age are shown in red and those that become hypomethylated with age in blue. (E-G) Bar plots showing comparisons between age-associated loci (dark gray) and background (light gray) coverage of genomic features. (H) Comparison of the change in methylation during early- and late-life across the top 1000 age associated loci. (I) Table showing differential methylation between early- and late-life. (J) Differences in the percent of reads which have discordant methylation across the three age groups.

### Genomic distribution of age-associated loci

To determine the organization of age-associated loci throughout the genome, the distribution of age-associated loci was mapped across chromosomes. Similar to findings in other species (10,15), age-associated changes appear to be widespread with both positively and negatively age-associated loci (n = 1000) present on all 24 chromosomes (Figure 1C). Significant age-associated loci (n=62) were also relatively evenly distributed across the genome with the exception of chromosomes 2, 3, 9, 14, 20 and 23 where we did not detect any CpGs with significant age-associations (Figure 1D). We then examined the genomic locations of the top 1000 age-associated CpGs, as determined by Spearman correlation coefficients, with respect to their proximity to CpG islands and genic context. We observed a tendency for age-associated sites to fall into non-coding regions with low CpG density. For example, age-associated loci are enriched in CpG shelves (*p* = 0.021) and open-seas (*p* < 1.1e-09), and are significantly depleted in CpG islands (*p* < 5.7e-16) which generally harbor gene promoters (39)(Figure 1E). Relative to background, a significant enrichment of age-associated CpGs located in introns (*p* < 5.3e-14) and intergenic regions (*p* < 0.0002) and a significant depletion of age-associated CpGs in exons (*p* < 2.2e-16; Figure 1F) was observed. CpGs in the human epigenetic clock are reported to co-localize with glucocorticoid response elements (23), so we tested for significant deviations from the expected overlap between age-associated sites and three hormone response elements. However, enrichment of age-associated CpGs in glucocorticoid, estrogen, an androgen response elements was not detected (Figure 1G).

### Temporal dynamics of epigenetic aging

Using the top 1000 age-associated loci, we determined that the mean absolute rate of change was greater in early life (2-6 months; 13.7%) when compared to later in adulthood (6-12 months; 9.7%; *p* < 2e-16; Figure 1H). After correcting for the number of days elapsed between each time point, medaka had an average absolute change in methylation of 0.11% per day during early life and 0.047% per day during later life, suggesting that the rate of change in methylation is more than twice as fast during early life. When sites at which age has the greatest effect size (greatest percent difference in methylation from 2-12 months; n = 1000) were analyzed, the rate of change was even more striking as the mean absolute methylation changes between 2- and 6-months is 22.1% compared to just 12.7% between 6- and 12-months (*p* < 2e-16). Additionally, the aging methylome also appears to be qualitatively more dynamic in early life, as 904 DMCs were observed between 2-6 month old individuals and only 352 DMCs observed between 6-12 month individuals (Figure 1I). Whereas the number of cytosines that become hypomethylated with age remain approximately constant in early and late time points, there are far more cytosines that become hypermethylated with age in early life (784 loci) compared to later life (236 loci), demonstrating that the difference between early and late life methylation is driven primarily by cytosines acquiring methylation with age (Figure 1I). Collectively, these findings demonstrate that the DNA methylome is both quantitatively and qualitatively more dynamic during early life.

### Discordant methylation

We also examined the relationship between chronological age and discordant methylation, an indicator of epigenomic instability (40). Genome wide levels of discordant DNA methylation were observed to increase with age (*p* < 2.1e-05; Figure 1J), suggesting that epigenetic patterns are indeed eroded over time consistent with epigenetic drift. Interestingly, differences in the proportion of discordant reads appeared greatest between 2- and 6-month old individuals, suggesting that both epigenetic drift and age-associated DNA methylation patterning are both more dynamic in younger fish.

### Constructing an epigenetic clock for medaka

Training an elastic net regularized regression model on the medaka DNA methylome revealed the highly predictable nature of age-associated remodeling. The modeling approach selected 39 CpGs which were collectively the most predictive of chronological age from a subset of 45,273 loci covered in all samples. The model performed well on the training set (cor = 0.99; mean error = 1.3 days) and was validated using a test set (n = 3) consisting of one sample from each age group that was selected at random (cor = 0.9495; mean error = 29.1 days; Figure 2). When considered in the context of a 2-year lifespan, which is standard in our medaka population, this represents age-prediction accuracy within 4% of the lifespan.

**Figure 2.**
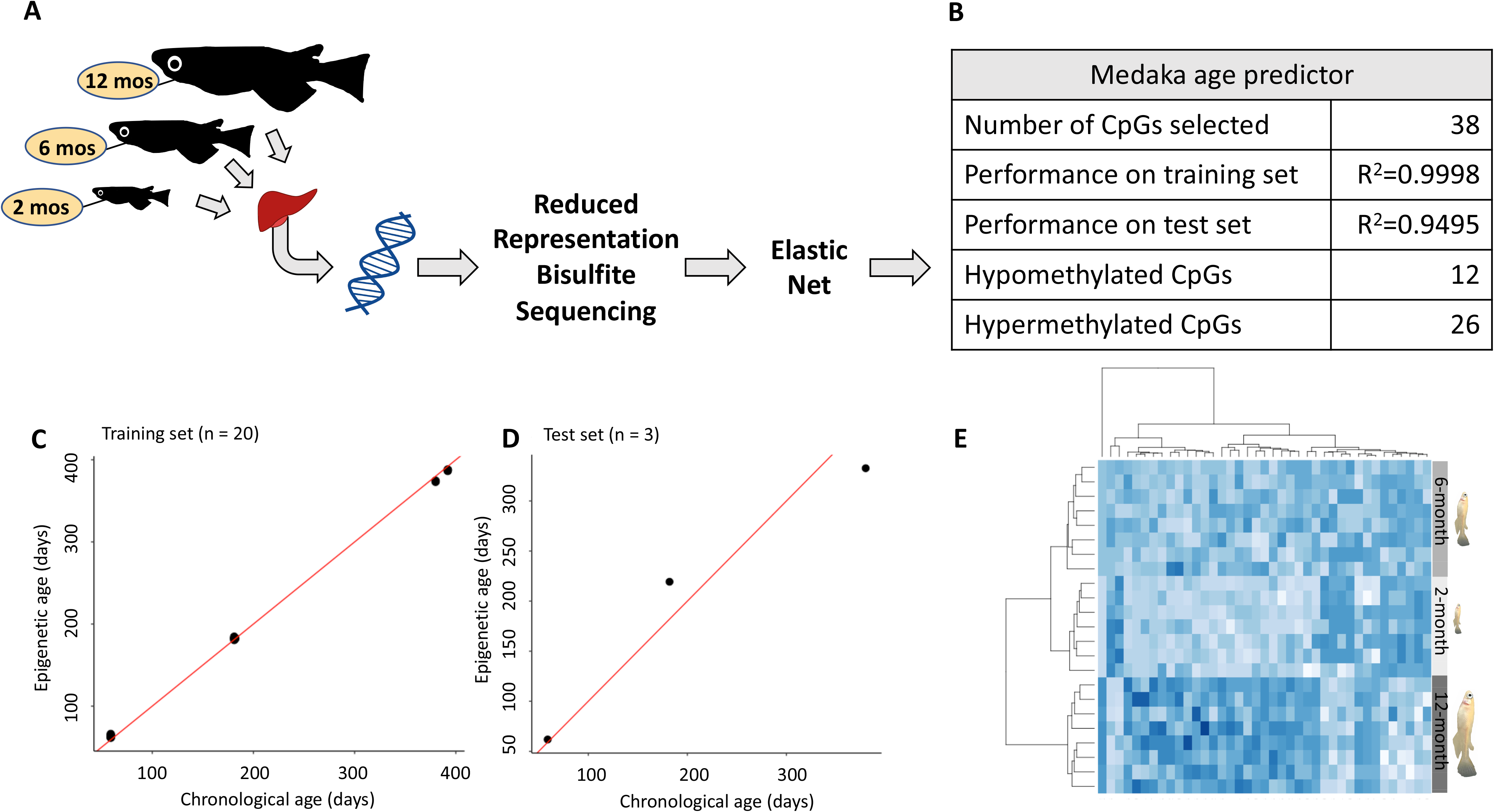
Construction of an epigenetic age predictor in medaka. (A) Conceptual diagram of the RRBS experiment. (B) Description and performance metrics of the medaka epigenetic clock. (C) Performance of the epigenetic clock on the training set (n=20) and (D) test set (n=3). (E) Heat map of the methylation of the 38 clock loci. Age is specified by color (2-month: light gray, 6-month: medium gray, 12-month: dark gray.)

### Environmentally accelerated epigenetic aging

To test if exposure to an environmental stressor accelerates epigenetic aging, we first built a new epigenetic clock using CpGs which were covered across all samples in both our initial age cohort and those exposed to ionizing radiation. Although this new clock incorporated a different set of CpGs than that of control fish alone, the mean error (37.0 days) was similar for the control individuals and the previous test set (29.1 days). When applied to fish exposed to ionizing radiation for seven weeks at 5, 50, and 500 mGy/day, exposed fish were on average 31.2 (13.3%), 41.3 (17.6%), and 32.4 (13.8%) days older than control individuals exposed to background levels of radiation, respectively (Figure 3A). However, given individual variability in epigenetic age estimates within exposure rates, this increase in epigenetic age was not statistically significant. Interestingly, IR exposure did not affect the percentage of reads with discordant methylation (Figure 3B), nor did it affect survival rate across treatments.

**Figure 3.**
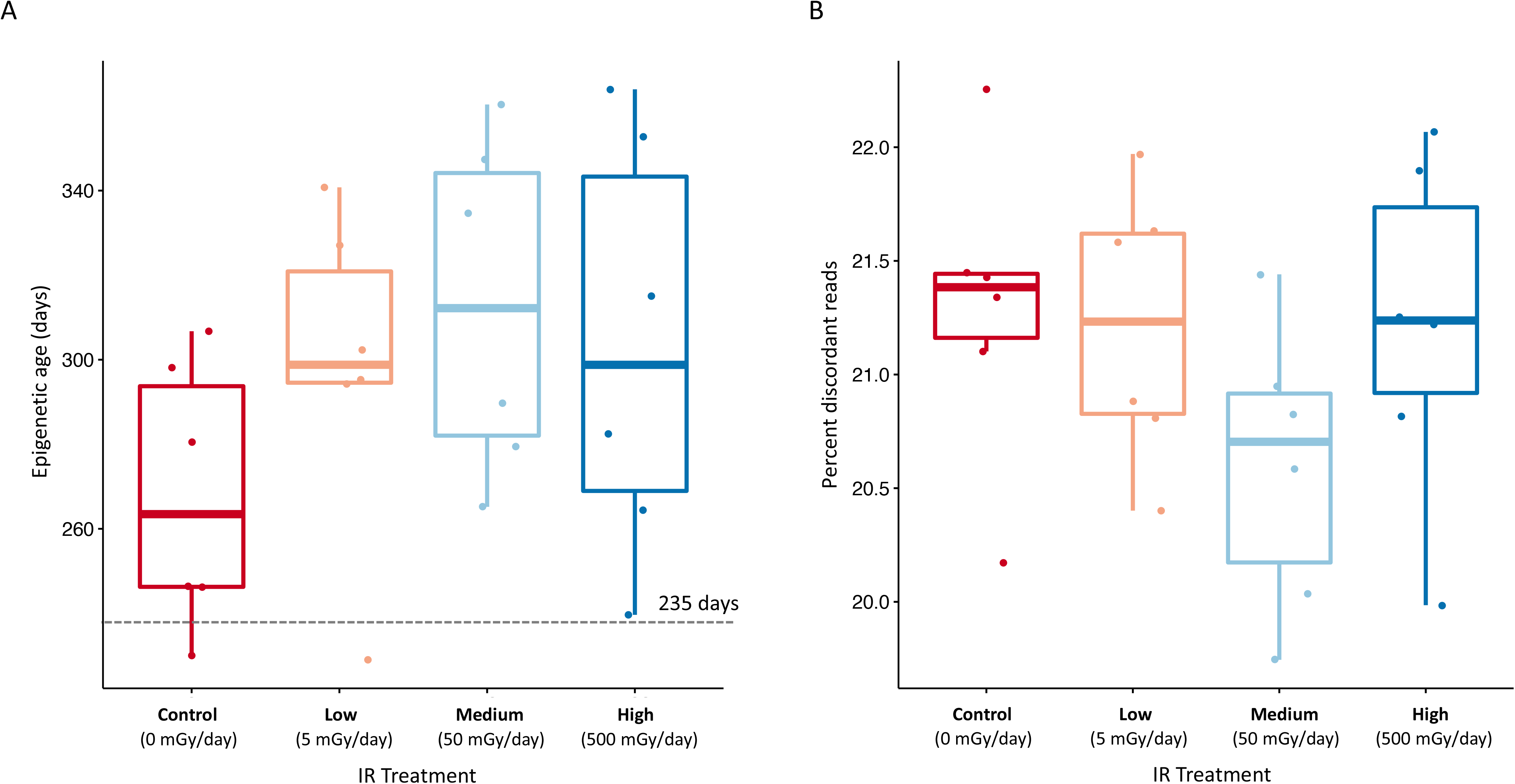
Effect of ionizing radiation on age-associated DNA methylation. (A) Epigenetic age predictions for fish exposed to 7-weeks of ionizing radiation at various dose rates (0, 5, 50, and 500 mGy/day). Average chronological age is shown by dotted gray line (235 days). (B) Percent of reads with discordant methylation across exposure groups.

### Environmental effects on the aging methylome

We hypothesized that the influence of age and environmental factors on CpG methylation might exist along an opposing continuum with strongly age-associated loci independent of environmental influences on one end and loci affected by the environment and less likely to be affected by age on the other. However, along this continuum, we posited that a subset loci exist that are affected both by age and environmental exposures (Figure 4A). To test this hypothesis, we first sorted CpGs into bins according to the degree to which their methylation status was correlated with age to form an aging continuum. We then determined which CpGs become differentially methylated after exposure to ionizing radiation and identified 3,909 loci whose methylation status is significantly affected by exposure to at least one dose of IR (5, 50, and 500 mGy/day) relative to our background exposure control. Of these differentially methylated cytosines, 666 (17% of IR DMCs) were also represented in our aging dataset (out of a total of 79,684 loci). We then tested if cytosines which were differentially methylated after IR exposure were enriched in age-associated bins. Whereas we hypothesized that these differentially methylated cytosines would be overrepresented in bins harboring loci with weak age-associations, we surprisingly found a two-fold enrichment in the bin containing loci which have strong positive correlations with age (cor = 0.5-1.0; Figure 4B).

**Figure 4.**
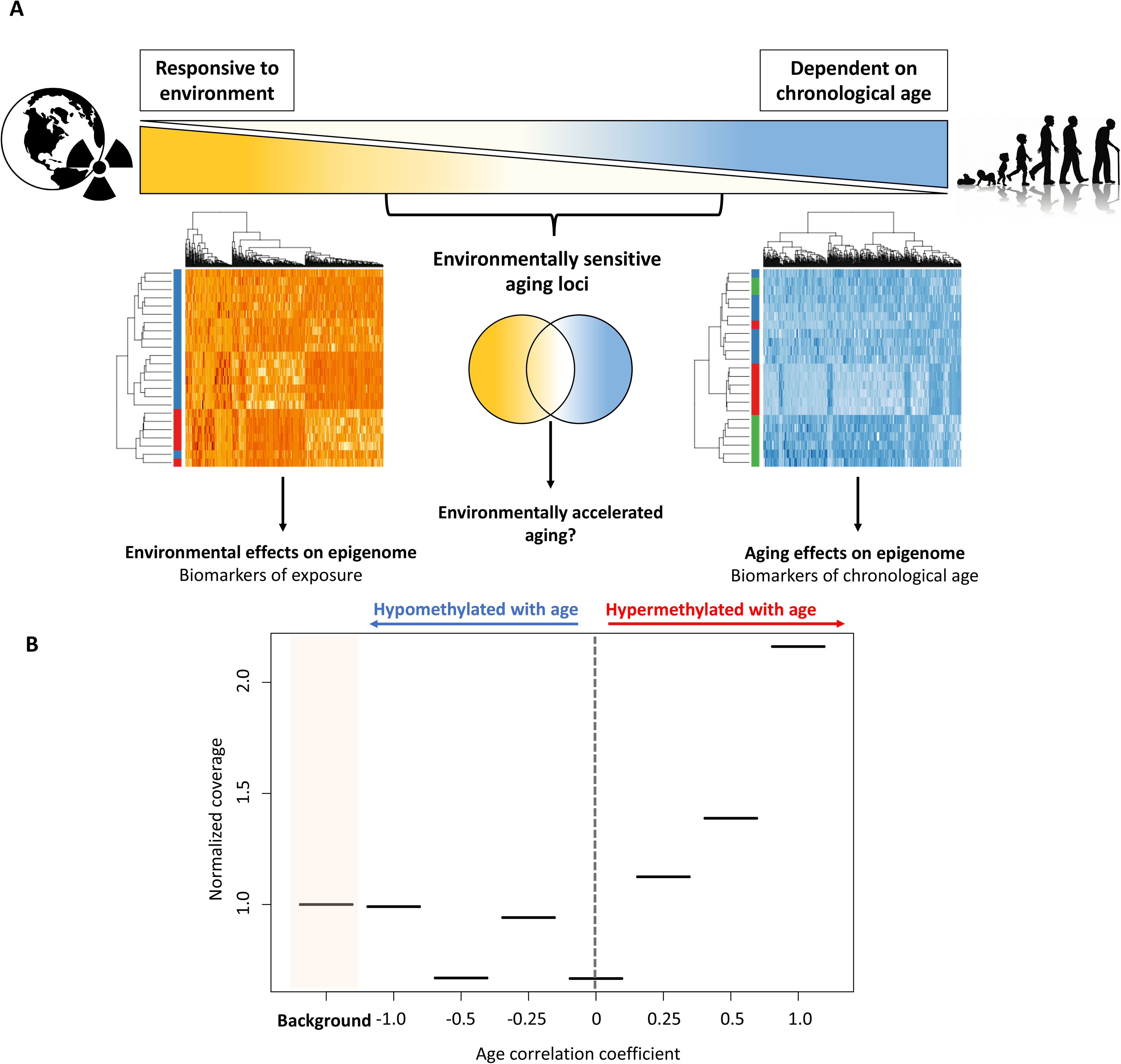
Interactive effects of exposure to ionizing radiation and age on the medaka DNA methylome. (A) Diagram of a hypothetical continuum of loci with loci affected strongly by age on one end and those which are highly environmentally responsive on the other. (B) Distributions of loci affected by IR exposure along the continuum of association with chronological age. Background represents the number of overlapping CpGs in the two datasets (age and radiation exposure).

We further investigated the link between exposure to IR and epigenetic aging by assessing the directionality of IR-induced methylation in relation to changes occurring during normal aging. Of the loci which were both strongly affected by IR and age (Figure 5A), we found that IR exposure resulted in methylation shifts reflective of both accelerated and decelerated epigenetic age (Figure 5A). Age-associated loci are split evenly between having IR-induced accelerated and decelerated patterns, and these effects are observed across the genome (Figure 5B). However, when the effects of IR exposure on the methylation status of strongly age-associated loci (absolute correlation coefficient >0.5; n = 33) were analyzed independently across IR dose, we observed dose dependent effects on the direction of change. Of these dually affected loci, 66.7% of loci affected by low doses of IR (5 mGy/day), and 83.3% of loci affected by medium doses of IR (50 mGy/day) shifted methylation patterns in a direction consistent with accelerate aging, whereas 73.3% of loci affected by exposures to high doses of IR (500 mGy/day) elicited changes in the opposite direction of normal epigenetic aging (Figure 5C). However, these dose dependent changes occur at different loci across the genome, and it is unclear if the difference in directionality is due to locus specific effects.

**Figure 5.**
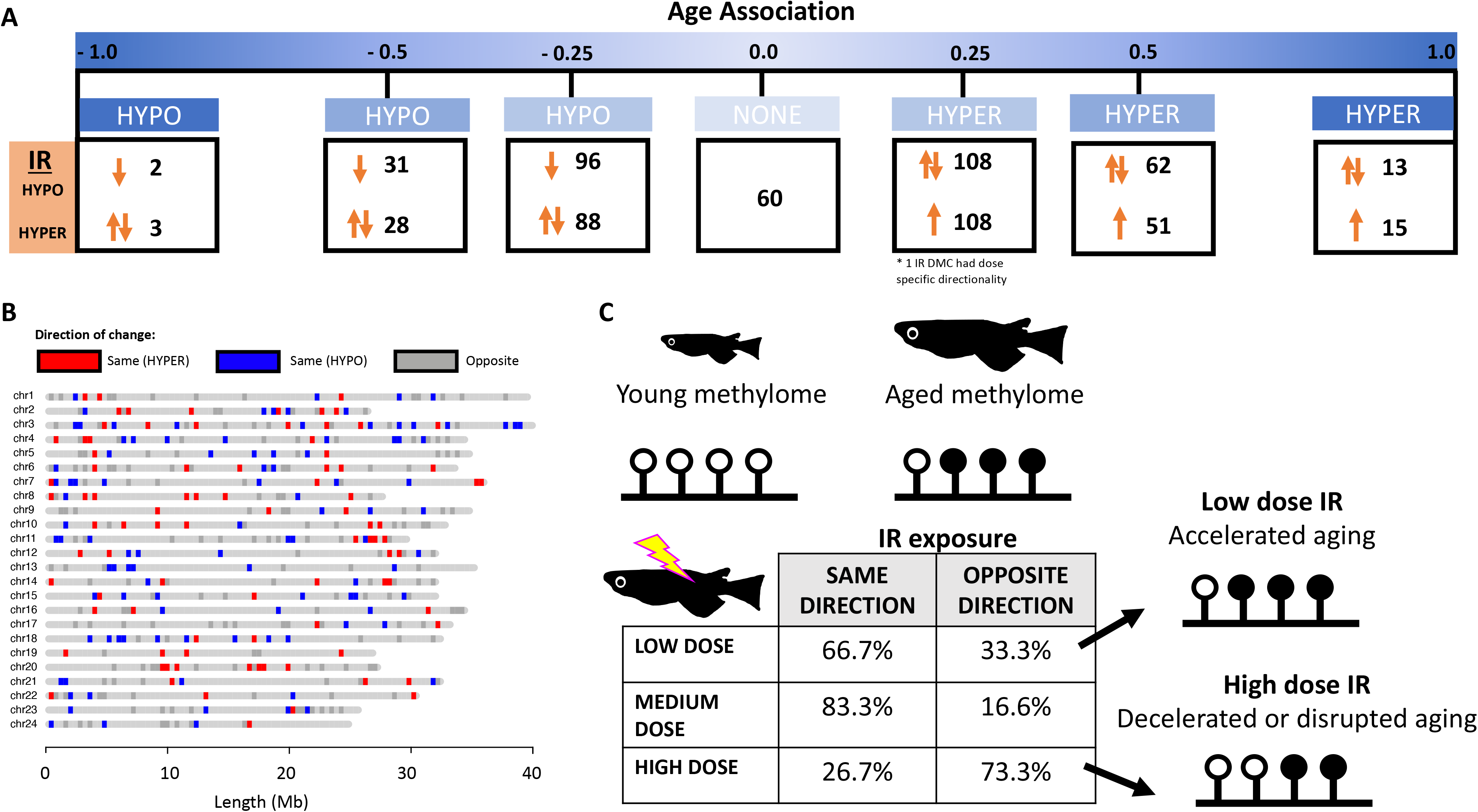
Directionality of IR-induced changes to methylation status in the context of normal epigenetic aging. (A) Distribution of loci which become differentially methylated from IR exposure along the continuum of association with chronological age. Arrows signify whether the IR induced change is in the same or opposite direction as changes induced by age. (B) Genomic distribution of loci which become differentially methylated with IR exposure in the same (red/blue) or opposite (gray) direction as age-related changes. Loci which become hypermethylated with both age and IR exposure are shown in red and those which become hypomethylated in blue. (C) Table showing the percent of loci whose methylation changes in the same and opposite direction across dose rates and a conceptual diagram of the hypothesized effect this could have on the aging methylome.

## Discussion

Here, we characterize age associated patterning of the medaka DNA methylome and model these patterns to construct an epigenetic clock that predicts chronological age with an accuracy of 4% of the lifespan. We demonstrate that epigenetic aging in medaka shares similarities with other species such as mice and humans (12). Most notably, age associated loci tend to be located in regions with low CpG density and in non-coding regions, similar to findings in mice (15). However, in contrast to humans, we do not observe an enrichment of age-associated methylation patterning in glucocorticoid receptor response elements, nor in the response elements of other nuclear hormone receptors, suggesting fundamental differences in aging programs are also present (23). How these similarities and differences translate into functional variation across aging and life history trajectories warrants further investigation.

Epigenetic aging in medaka occurs approximately twice as fast early in life when compared to rates in mature fish. This is consistent with findings in mice, humans, and dogs suggesting that the methylome is most dynamic early in life and corresponds to physiological processes (i.e. growth, maturation) in similar ways across species (21). In addition to increased rates of change at specific CpGs, we find that a greater number of loci incur methylation remodeling during early life. These findings raise the potential that epigenetic aging occurs on a qualitative scale in which different loci acquire or lose age-associated methylation patterning semi-independently of aging rate. Our data and those of others (2,41) are consistent with studies demonstrating that biological aging, as measured by telomere length, is also more dynamic during early life (42–46). In most cases, increased rates of telomere attrition are associated with faster growth early in life and are thought to reflect trade-offs between growth and longevity (47,48). Whereas positive correlations between accelerated epigenetic aging and individual variation in body size have been reported in humans (49), these linkages have not been explored in the context of variable life history strategies present in other ecological systems (12). Given that evolutionary theories of aging, such as antagonistic pleiotropy, emphasize the importance of maximizing fitness during early life (50,51), understanding the role of the environment in determining rates of epigenetic aging at different life stages stands to better inform our understanding of the mechanisms driving adaptive plasticity in life histories. Studies examining how epigenetic aging trajectories, both in a quantitative (e.g., rate) and qualitative (e.g., variable subsets of loci acquiring age-associated patterns) sense, are affected in early life are clearly needed to advance our understanding regarding both the origins of epigenetic-to-chronological age discordance and the potential adaptive roles it might play in nature.

Double stranded DNA breaks have recently been shown to accelerate epigenetic aging in mice (37), and based on these findings, we hypothesized that chronic IR exposure, a known source of DSBs (52), might also result in accelerated aging. Previous research in humans has demonstrated a relationship between radiation treatment in breast cancer patients and increased epigenetic age (53). However, after a 7-week exposure to three doses of IR, we only observe a trend towards accelerated epigenetic aging in medaka characterized by high individual variation within treatment groups. However, on average, IR resulted in an average acceleration of 35 days across all exposure groups, which represents a 10-20% increase in age relative to controls. Given that IR is known to affect age-related phenotypes (54), such as increasing rates of telomere attrition (55), decreasing regenerative capacity (56), altering age-associated gene expression patterns (57), and increasing the accumulation of age-associated proteins (56), it is possible that our trend towards increased epigenetic age is another measure of accelerated aging induced by IR. Yet, whether our results reflect a weakly trained epigenetic clock, low sample size, or the unexpectedly high variation in individual age estimates is not clear. It is also possible that the timing of IR exposure dictates the impact of on epigenetic aging trajectories. The fish in this study were approximately 6-months old at the start of the IR exposure and given that the rate of epigenetic aging slows with chronological age (2), exposure during earlier windows of development are likely to be more influential. Experiments assessing different exposure durations at different life stages are needed to determine if environmentally relevant sources of DSBs result in epigenetic-to-chronological age discordance.

Although the epigenetic clock itself was not affected, IR exposure did interact with the aging DNA methylome in unexpected ways. CpGs which become strongly hypermethylated with age were more likely to be affected by IR exposure. This result ran counter to expectations as we hypothesized that loci strongly affected by age would be relatively insensitive to environmental factors. Interestingly, most of the methylation changes occurring in early life were due to sites which become hypermethylated and are thus more likely to be affected by IR exposure, which adds further support to the idea that environmental exposures in younger individuals might disproportionately impact epigenetic age acceleration. We also found that the directionality of IR-induced shifts to the methylation status of age-associated loci was dose-dependent with exposure to low doses of IR resulting in patterns consistent with precocious aging and high doses inducing shifts towards younger states. This could in part be why we see large variation in the age prediction of higher dose rates. IR has long been considered a general accelerant of normal aging (54), and yet our findings suggest the effects of IR on the aging epigenome are more nuanced than previously realized and may both accelerate and decelerate normal patterns of epigenetic aging. IR exposure, especially at relatively low doses over long time scales, is capable of eliciting variable outcomes (58,59), and the response of epigenetic aging trajectories to these ecologically (and evolutionarily) relevant exposures will likely inform a broader understanding of the potentially adaptive role of epigenetic age acceleration in nature.

The current study demonstrates that specific aspects of epigenetic aging are broadly conserved across vertebrates, while other aspects appear more divergent. Given the similarities between the medaka epigenetic clock with those developed in mammals, epigenetic aging is likely to have a conserved functional role in the aging process, although more work examining the causal roles of epigenetic aging is needed. Further, using these predictable patterns we have developed a medaka epigenetic clock which can be used to predict chronological age as well as measure biological age acceleration after exposure to environmental stressors. Our results demonstrate that the timing of exposure may determine the strength of those relationships, however this has been relatively unexplored. As we report a strong dependence of epigenetic aging patterns on early life changes, the time period prior to sexual maturity may be particularly sensitive to environmental stressors. We also observe dose-specific effects of IR which alter the methylome in ways that both accelerate and decelerate normal epigenetic aging. These findings nicely demonstrate how the high resolution associated with DNA methylome sequencing adds nuance to our understanding regarding the environmental drivers of biological aging. Overall, this study provides insight into epigenetic aging processes in vertebrates and establishes a model by which environment-by-aging interactions can be further explored.

## Methods

### Development of the epigenetic clock

#### Animal husbandry and rearing

All procedures involving fish husbandry and animal experiments were approved by the University of Georgia’s IACUC (protocol A2018 09-007-Y3-A2). Medaka were bred under optimal conditions (24°C, 16L:8D) to produce offspring which were raised under similar conditions. For the construction of the epigenetic clock, male fish aged 2-months (n = 7), 6-months (n = 8), and 12-months (n =8) were sacrificed using an overdose of sodium bicarbonate buffered Tricaine (MS-222 300 mg/L). Fish were immediately necropsied, and hepatic tissue stored in RNAlater at −80°C. The sex of fish was determined using sexually dimorphic fin structure and gross morphology of gonads. Animals at 2 months of age had not yet reached sexual maturity so PCR amplification of the Y-chromosome specific gene, dmy, was used to determine genetic sex (60).

### DNA extraction

DNA was extracted using a modified column protocol. Briefly, whole livers were homogenized in a Mini-Beadbeater (BioSpec, Bartlesville, OK) using stainless steel beads for 2 minutes at 30 Hz in lysis buffer (4M guanidinium thiocyanate, 0.01M Tris-hydrochloric acid (pH 7.5), and 2% beta-mercaptoethanol). Resulting lysates were centrifuged, and supernatants were transferred to spin columns containing a fiberglass filter (Epoch Life Science, Missouri City, TX). Column-bound DNA was washed twice with wash buffer (60% ethanol, potassium acetate (162.8 mM)), and tris-hydrochloric acid (27.1 mM, pH 7.5), diluted with 60% (v/v) of ethanol) and eluted in 40 μl of TE buffer. DNA concentrations were quantified using a Qubit fluorometer 2.0 (Invitrogen, Carlsbad, CA) and purity was assessed on a Nanodrop spectrometer (Thermo-Scientific, Waltham, MA) using absorbance ratios at 260/280 and 260/230. DNA was stored at −20°C until library preparation.

### Reduced Representation Bisulfite Sequencing

Reduced representation bisulfite sequencing (RRBS) was used to analyze the DNA methylome, which is preferred over whole genome bisulfite sequencing due to the enrichment of areas with high CpG content (61). The RRBS libraries were prepared using the Diagenode Premium RRBS Kit (#C02030032, Diagenode, Denville, NJ) with the following exceptions: 200 ng of genomic DNA was used for input and the equation cycle threshold (Ct) + 2 was used to determine the number of PCR cycles needed for final library amplification. AMPure XP Beads (Beckman Coulter, Brea, CA) were used for all steps in the Diagenode protocol requiring size selection or clean-up of products. Final libraries were stored at −80°C until sequencing.

### Sequencing, Quality Control and Bisulfite Conversion Efficiency

Libraries were submitted to the Georgia Genomics and Bioinformatics Core where they were further assessed for quality and concentration using a Fragment Analyzer (Agilent, Santa Clara, CA). Libraries were pooled and sequenced single-end for 75 cycles on one high-output flow cell of an Illumina NextSeq with 5% PhiX control (Illumina) added to provide sequence diversity. The sequencing run generated 225 million reads across samples, with 92% of reads falling above the high-quality threshold of Phred score >30. Individual samples had between 4.4-9.9 million reads and their quality was assessed using FastQC (v0.11.5). Adapters and low-quality sequences (Phred score <25) were trimmed using TrimGalore! (v 0.4.5; --rrbs) and 4bp at the 3’ end was removed due to low overall quality scores of the 3’ end of reads. The efficiency of the bisulfite conversion was determined to be >98%.

### Alignment and MethylKit

Using Bismark (v0.20.0) (62) the medaka reference genome (ASM223467v1) was indexed for bisulfite conversion, and reads were aligned to this indexed reference using Bismark with an overall alignment rate of 50-60%. Using the Bismark methylation extractor, BAM files containing methylation calls for each sample were produced, sorted using SAMtools (v 0.1.19)(63), and used as input for Bioconductor’s (v 3.9) package MethyKit (v1.10.0) (64) in Program R (v3.5.1). In MethylKit, we first filtered out CpGs which were covered at a depth of less than 2x or greater than 100x and normalized samples by coverage using MethylKit’s normalizeCoverage function to prevent bias arising from PCR or over-sampling of specific individuals, respectively. For characterization of the aging methylome, CpGs which were covered at a depth of at least five reads in at least 90% of samples were selected resulting in a subset of 79,684 CpGs which were used for further analysis. A reduced subset of 45,273 CpGs, which were covered in all samples at a depth of at least 5 reads, was selected for the development of the epigenetic age predictor using an elastic net penalized regression model which requires no missing data.

### Correlation between age and methylation status

We used Program R (v3.5.1) and the function corr.test from package psych (v1.8.12) to run 79,684 independent Spearman correlations for each CpG which passed filtering. Correlation coefficients between the percent methylation and age in days were computed and significance values were calculated using a false discovery rate (FDR) correction for multiple comparisons.

### Determination of differentially methylated cytosines

To determine differentially methylated cytosines we used MethylKit’s (v1.10.0) ‘calculateDiffMeth’ function. Differentially methylated cytosines were those which had at least 25% difference in methylation between age groups with a q-value less than or equal to 0.01. Three individual comparisons were made between age groups (2-month vs. 6-month, 6-month vs. 12-month, and 2-month vs. 12-month.)

### Characterization of Genomic Location

Genomic locations for age-associated loci determined by Spearman correlations were characterized using Galaxy’s coverage function (Galaxy Version 2.29.0) (65). To determine the genic context of each CpG, we used the medaka reference genome annotation (ASM223467v1) to assign CpGs as lying in introns, exons, or intergenic (not in exon or intron) regions. Similarly, CpG island (CGI) context was determined by first producing a BED file with CGI coordinates using Galaxy’s function cpgplot (Galaxy Version 5.0.0) (65) according to a sliding window of 100 bp (minimum size of CGI = 200 bp, Minimum average observed to expected ratio = 0.6, Minimum average percentage of G plus C = 50.0). Using bedtools ClosestBed (Galaxy Version 2.29.0) (65), we were then able to assign each CpG to either a CpG island (within CGI), CpG shore (area of 2kb flanking CGI), CpG shelf (area from 2kb-4kb flanking CGI), or as an open-sea CpG (further than 4kb to nearest CGI). If the CpG fell within multiple categories due to being proximate to more than one CGI, ties were reported and the CpG was counted in both categories (n = 4). Enrichment above background was performed with binomial tests using the full dataset of 79,864 covered loci as the background.

### Hormone response elements (HRE)

To determine if age-associated loci were enriched in hormone response elements (HREs), we first established the coordinates of prospective HREs in the medaka genome using PoSSuM search (Version 2.0). Raw PFM files were downloaded from JASPAR (66) for the estrogen (MAO112.2), glucocorticoid (MAO113.3), and androgen response elements (MA0007.2). The raw PFM files were transformed into PPM files (divide the frequency by the total count for that position) and then into PSSM files (Log2(PPM value/0.25)) where the PPM value is the count/total and the 0.25 is the null expected frequency of that base. These PSSM files were input into PoSSum search (-pr -format tabs -pval 0.0001 -lazy -uniform -lahead -dna) and compared against the medaka reference genome to identify prospective HREs with a p-value < 0.0001 considered a significant HRE. The genomic coordinates from the prospective HRE sites were extracted into a BED file and compared against the age-associated loci and background loci using the coverage function in Galaxy (version 20.05) (65) to determine coverage in a 2kb window surrounding each CpG (1kb upstream, 1kb downstream). Enrichment above background was performed using binomial tests using the full dataset of 79,864 covered loci as the background.

### Early vs late rate of change

Using 1000 loci with the greatest Spearman correlations with age, we calculated the average absolute rate of change in methylation (absolute[% methylation late - % methylation early]) between each of the age groups (2- to 6-months, 6- to 12-months). Due to the differences in the amount of time elapsed between early (2- to 6-months) and late (6- to 12-months) time points, we normalized the average absolute rate of change by number of days between each age interval. This ‘daily’ average absolute rate of change was compared between the early and late time points using a t-test.

### Elastic net penalized regression model

To build the epigenetic age predictor, we used the GLMNET package (v1.9-9) (67) in Program R (v3.5.1). This approach is similar to those used for epigenetic clocks developed in human (10) and mouse models (15). We used an elastic net penalized regression model (alpha = 0.5, family = gaussian) to select CpGs and assign penalties to individual model coefficients using the subset of 45,273 loci covered in all samples at a depth of ≥ 5x. We used a 10-fold cross validation to select the optimal lambda value (value resulting in minimum mean error) for the model on the training set (n = 20 samples) and was verified on a test set (n = 3; one sample from each age group) that was selected at random.

### Testing the epigenetic clock

#### Age prediction in fish exposed to ionizing radiation

Six month old male fish were exposed to low doses of ionizing radiation for 7 weeks at the Savannah River Ecology Laboratory’s Low Dose Irradiation Facility (LoDIF) (68). The irradiation protocol is outlined in Bertucci et al. (2020) with slight modifications (69). Briefly, eggs were collected during a 14-day window in January and February 2019 and reared in mixed-sex cohorts as described above. Males were separated and put into 19 L containers (15 fish/container) and placed under irradiators equipped with Cesium-137 sources emitting radiation at approximately 5, 50 and 500 mGy/day (69). During the exposure period, fish were fed three times per week with Tetramin and were given a fresh flow of water for an hour at each feeding. Radiation sources were turned off briefly during feeding. After a 7-week exposure, fish were immediately euthanized with an overdose (300 mg/L) of Tricaine/MS-222 and immediately necropsied. DNA from hepatic tissue (n = 23) was isolated and library preparation for RRBS was performed as described above. Sequencing was performed at the GGBC as described above, with the addition of 20% Illumina PhiX control spiked in (for greater base pair diversity.) Sequencing resulted in 300,267 CpGs which were covered across all samples at a depth of at least 5 reads. To assess epigenetic age in IR exposed individuals, CpGs covered across all samples from the original aging cohort and this experiment were identified. We created an epigenetic age predictor by training it on all 23 samples from the aging cohort and tested it on the 24 samples from the LoDIF experiment. Epigenetic age predictions were compared between groups using analysis of variance tests (ANOVA).

#### Determination of IR Differentially Methylated Cytosines

MethylKit’s (v1.10.0) ‘calculateDiffMeth’ function was used to determine differentially methylated cytosines between control and exposure groups. Differentially methylated cytosines (DMCs) were those which had at least 25% difference in methylation between exposure groups with a q-value less than or equal to 0.01. Three individual comparisons were made between control and treatment groups: Control vs Low (5 myGy/day), Control vs Medium (50 mGy/day), and Control vs High (500 mGy/day).

#### Overlap of age-associated sites with IR DMCs

The overlap between the age-associated loci (as determined by Spearman correlations) and the IR DMCs was analyzed by creating BED files with the coordinates of each loci and using Galaxy’s coverage function (Galaxy Version 1.0.0) (65) to determine the overlap between the age-associated and IR altered loci.

#### Percentage of discordant reads

The percentage of discordant reads (PDR) was calculated using the ‘methylation_consistency’ extension in Bismark (v0.20.0, Babraham Bioinformatics). First, the number of reads containing at least 5 CpGs which had concordant (0-9% or 91-100% methylated) or discordant (10-90% methylated) patterns of DNA methylation was determined. Then the PDR was calculated as the number of discordant reads divided by the total number of reads with at least 5 CpGs covering that locus.

## Abbreviations

DMC: Differentially methylated cytosine
DNAm: DNA methylation
FDR: false discovery rate
IR: Ionizing radiation
LoDIF: low dose irradiation facility
PCR: polymerase chain reaction
PDR: Percentage of discordant reads
qPCR: quantitative polymerase chain reaction

## Acknowledgments

The authors would like to dedicate this work to the memory of James Taylor, one of the visionaries and founders of The Galaxy Project. The authors also acknowledge Samantha Bock, Matthew Hale, and Faith Leri for helpful discussions in the design and analysis of this research. The authors would also like to acknowledge and thank Donald Mosser, Mark Edwards and the SREL maintenance staff, and Matt Wilmot and the DOE radiation control staff for help in logistics in safely operating the LoDIF.

## Disclaimer

This report was prepared as an account of work sponsored by an agency of the United States Government. Neither the United States Government nor any agency thereof, nor any of their employees, makes any warranty, express or implied, or assumes any legal liability or responsibility for the accuracy, completeness, or usefulness of any information, apparatus, product, or process disclosed, or represents that its use would not infringe privately owned rights. Reference herein to any specific commercial product, process, or service by trade name, trademark, manufacturer, or otherwise does not necessarily constitute or imply its endorsement, recommendation, or favoring by the United States Government or any agency thereof. The views and opinions of authors expressed herein do not necessarily state or reflect those of the United States Government or any agency thereof.

